# Shift from a Simplified to Complex Gut Microbiota Reduces Adenoma Burden in a Preclinical Rat Model of Colon Cancer

**DOI:** 10.1101/2023.01.20.524931

**Authors:** Susheel Bhanu Busi, Daniel Davis, Jacob Moskowitz, James Amos-Landgraf

## Abstract

Specific bacterial taxa in the gastrointestinal tract have been strongly associated with cases of colorectal cancer (CRC) cancer in familial adenomatous polyposis and spontaneous disease cases in humans. This has been recapitulated in animal models of CRC with positive correlations with many commensals and pathogens. However, many of these studies are performed either in germ-free animals or employ an antibiotic regime, overlooking the complex interactions of the commensals within the colon. To simplify the challenges associated with the complexity of the microbiota in the GI tract we established the Pirc rat model of colon cancer on an Altered Schaedler Flora (ASF) gut microbiota (GM), maintained in a barrier room. To elucidate the role of the simplified (ASF) and conventional GMs on disease susceptibility, We conventionalized ASF Pirc littermates. We found that the conventionalized F1 rats had increased microbial diversity and decreased colonic adenoma multiplicity. Our findings show that the complexity and the interactions of the GM community and not a *Firmicutes* to *Bacteroidetes* ratio are an important factor affecting disease susceptibility.

## Introduction

Colorectal cancer (CRC) models, including both mice and rats, have been used to understanding the etiology of human diseases for decades ^1,2^. The ideal model should recapitulate the phenotype observed in humans, but also elucidate contributing factors such as the host microbiota and its relationship to the mechanisms of the disease. Recent evidence suggests that the gut microbiome, i.e. the collection of microorganisms in the large intestine plays an important role in the etiology of the disease ^3,4^. Several studies have tried to elucidate the mechanisms by which specific bacteria contribute to disease susceptibility by various methods including the utilization of germ-free ^5,6^ or monocolonized animals ^7,8^, or the use of antibiotics to eliminate endogenous gut microbiota (GM) populations ^3^. These majority of studies use the *Apc^+/Min^* mouse model that develop the majority of their tumors in the small intestine unlike human disease. Since the GM population has been shown to be different in the small intestine compared to the colon the translatability of these studies may be limited.

The Pirc (F344/NTac-*Apc^+/am1137^*) rat model of human colon cancer demonstrates a more consistent colonic tumor phenotype compared to the *Apc^+/Min^* mice and has been shown to have an altered phenotype with altered gut microbiota ^2,9,10^. To model more closely, the large number of endogenous commensals found in human CRC patients, we previously showed that the endogenous GM could be modulated through complex microbiota targeted rederivation ^2,9–11^. Determining the mechanisms and most importantly the interactions between commensals still pose challenges, considering the multiple permutations and combinations with the taxa found in the model.

In order to tackle the challenge of complexity, we established the Pirc rat on an Altered Schaedler Flora (ASF) gut microbiota ^12^. The ASF model includes an eight-community gut microbiome derived from mice, was developed in 1970 by R.P. Orcutt ^13^. Considering the highly diverse microbiome in animal models, the ASF community allows for in-depth identification of host-microbe interactions, including their spatial arrangment ^12^. Instituting Pirc rats on a minimal GM profile could potentially serve as a model for understanding mechanisms and interactions of specific bacteria, in the context of a well-defined, yet complex gut microbiome profile. Using CRASF (Charles-River ASF) rats as surrogates, F1-Pirc rats were established, and at weaning, littermates were transferred from a barrier room to a conventional status room in the animal facility. We hypothesized that transferring the Pirc rats to a conventional room compared to the cleaner, barrier room would increase the colonic tumor burden at sacrifice. Contrary to our hypothesis, we found that the animals maintained in the barrier (clean) room had significantly more colonic adenomas. This is the first time Pirc rats have been established on an Altered Schaedler Flora gut microbiota, but more importantly, suggest an even more central role for the gut microbiota in modulating the colon tumor phenotype of animal models for studying human diseases.

## Results

### Nominal taxa incursion in the Charles River Altered Schaedler Flora (CRASF)

In order to establish F344/NTac-*Apc^+/am1137^* rats onto a CRASF gut microbiota (GM), we first had to ensure that the simplified GM profile could be maintained in our facility. We housed four female and four male LEW/Crl ASF (CRASF) rats in a barrier room setting with individually ventilated racks in microisolator cages. Fecal samples collected prior to arrival at the facility and upon housing for 3 months at the facility, showed minimal addition of species to the GM profile. Over time, the LEW/Crl ASF animals acquired *Lachnospiraceae UCG-001, Lachnospiraceae UCG-006, Anaerotruncus, [Eubacterium], Enterococcus and Staphylococcus* (Fig.1a). Several cross-fostering experiments were needed to obtain a Pirc rat on a CRASF GM background. Out of 4 that were successfully cross-fostered, we only had one Pirc male rat. The F344/NTac-*Apc^+/am1137^* rat that was so cross-fostered onto the CRASF surrogates, at 1 month of age, showed a stable GM similar to that of the CRASF rats (Fig.1a). The ZymoBIOMICS mock microbial community standards simultaneously only acquired *Enterobacteriaceae*, potentially via the sample processing or sequencing or the bioinformatics analysis and annotation pipeline. Interestingly, the incursion of six taxa into CRASF rats led to significant differences when visualized using a Principal Coordinate analysis (PCoA) to understand the similarities between samples pre- and post-arrival, using the bray-curtis distance matrix (Fig.1b). The majority of the GM was maintained stably after housing the CRASF animals in a barrier setting for 3 months. To determine if the OTUs were acquired as a means of shipping to our facility, sequencing was performed on the bedding and gel-paks that the animals arrived with and found that four of the OTUs were possibly assimilated through the gel-paks, with *Muribaculaceae* making a significant contribution to the overall GM profile (Fig.1c).

**Figure 1.**
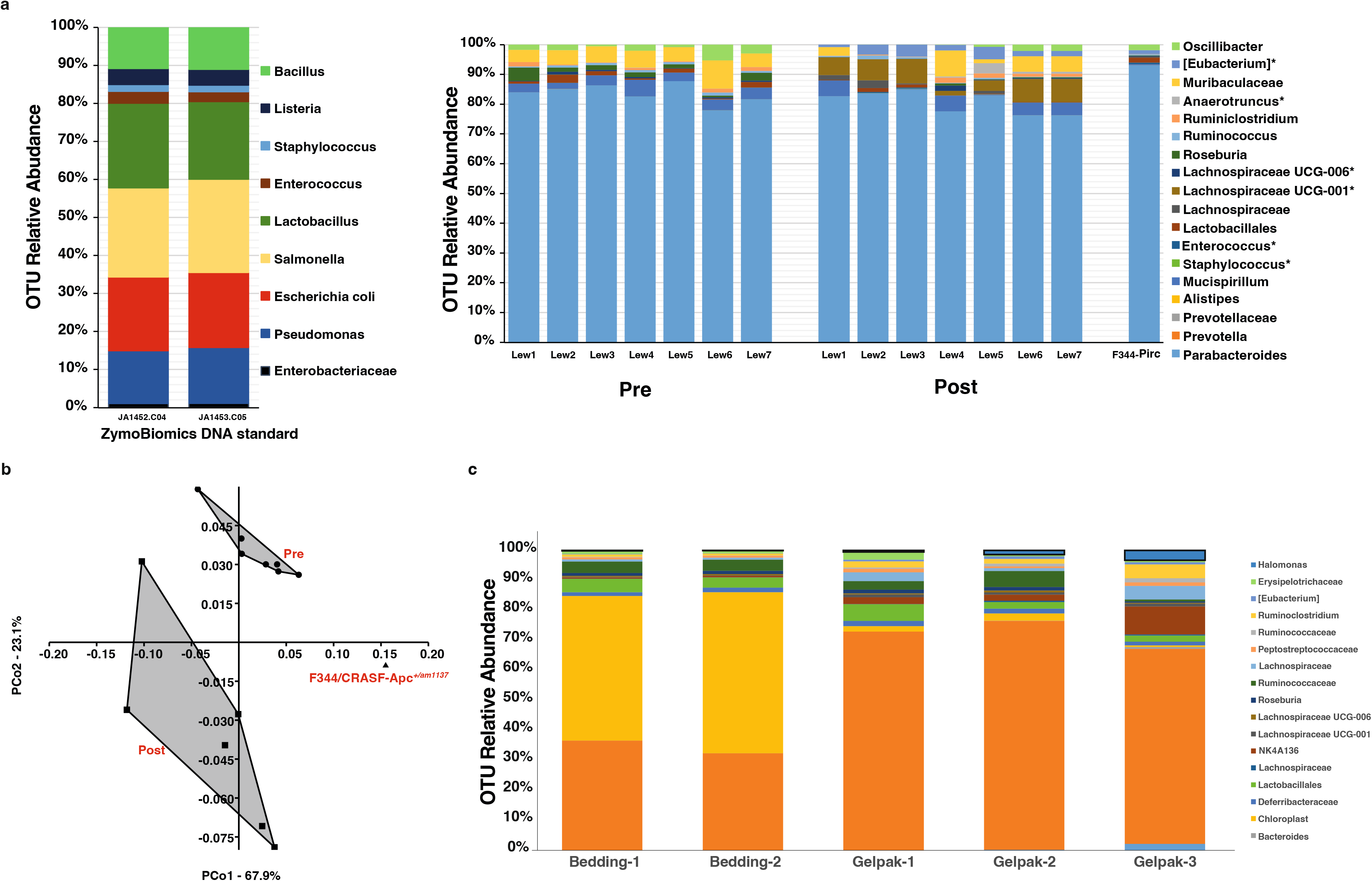
16S sequencing analysis of fecal microbiota in CRASF rats pre and post shipping. (a) Relative abundance (percentages) of each operational taxonomic unit (OTU) at the Genus level is shown for the ASF rats purchased from Charles River Laboratories, before shipping and 3-months post arrival at the Discovery Ridge animal facility. Also shown is the GM profile of the F344/CRASF-*Apc^+/am1137^* (JA1047.D4) that was fostered onto a CRASF dam. Bar graphs depicting the 16S sequencing data for the ZymoBIOMICS™ microbial community DNA standard is shown on the left that were used as processing and sequencing controls. *OTUs picked up after arrival and housing for 3 months. Lewis rats 1-4 were females, while rats 5-7 were males. (b) Principal Coordinate Analysis (PCoA) for the 16S rRNA sequencing data shows that Pre and Post samples (black, filled circles) of the CRASF rats are significantly different (PERMANOVA, F=6.272 and *p*=0.0001). The fostered *Apc^+/am1137^* rat is shown as the purple filled triangle. (c) Bar graphs representing each OTU as a single color show the relative abundance of taxa detected in the bedding and the gel-paks via 16S rRNA gene sequencing.

### Simplified gut microbiota increases susceptibility to colonic adenomas

LEWF344F1-*Apc^+/am1137^* CRASF rats obtained via the breeding set up were used to understand how the complexity of the gut microbiota (GM) may modulate disease susceptibility to adenomas in the rat model of colon cancer. At weaning, F1 Pirc littermates were separated into two separate rooms of the animal facility; a barrier room, where all cage changes were performed in a biocontainment hood, and a conventional room (Supplementary Fig.1). We found that the animals housed in the conventional setting had significantly fewer colonic adenomas than those housed under barrier conditions (Fig.2a). This differential tumor abundance was found in both male and female F1-Pirc rats. Interestingly, male rats from the conventional room had significantly more small intestinal tumors compared to the barrier rats, while female rats showed a similar trend (Fig.2b).

**Figure 2.**
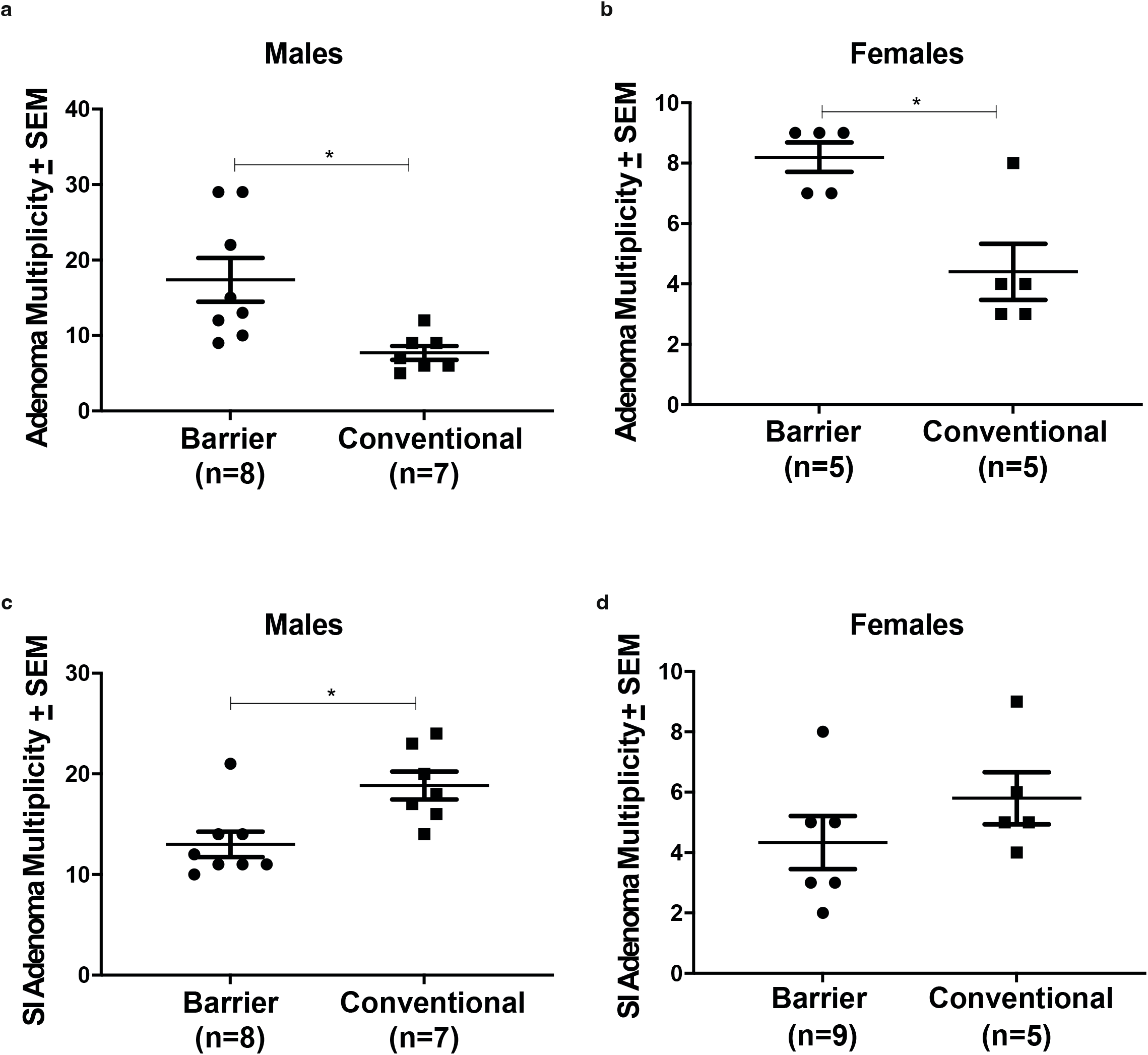
Colonic and small intestinal adenoma multiplicity of barrier and conventional rats at 4 months of age. Colonic (a) and small intestinal (b) adenoma multiplicity for male and female F1 Pirc rats from the barrier and conventional rooms is shown with adenoma counts on the y-axis and the groups on the x-axis. Significance was assessed by a Student’s *t-*test, with a *p-*value less than 0.05 was observed. Error bars indicate standard error of the mean (±SEM).

### Altered Schaedler Flora alters the colonic adenoma phenotype and the physiology of the gastrointestinal tract

Animals housed in the barrier room post-weaning demonstrated an increase in the number of proximal adenomas compared to conventionalized CRASF Pirc rats (Fig.3a). Most of these adenomas were 1 mm or smaller in diameter, however the rats with conventional GM did not show a similar phenotype (Fig.3b). Only one of twelve F1 Pirc rats separated at weaning and housed under a conventional settings had adenomas in the proximal colon that was slightly larger than 1 mm (Table.1). We also found that the overall number of small adenomas was significantly higher in the barrier room animals, irrespective of sex (Fig.3c), and the adenomas larger than 1 mm did not show any significant differences between the barrier and conventional rats’ (Fig.3d).

**Figure 3.**
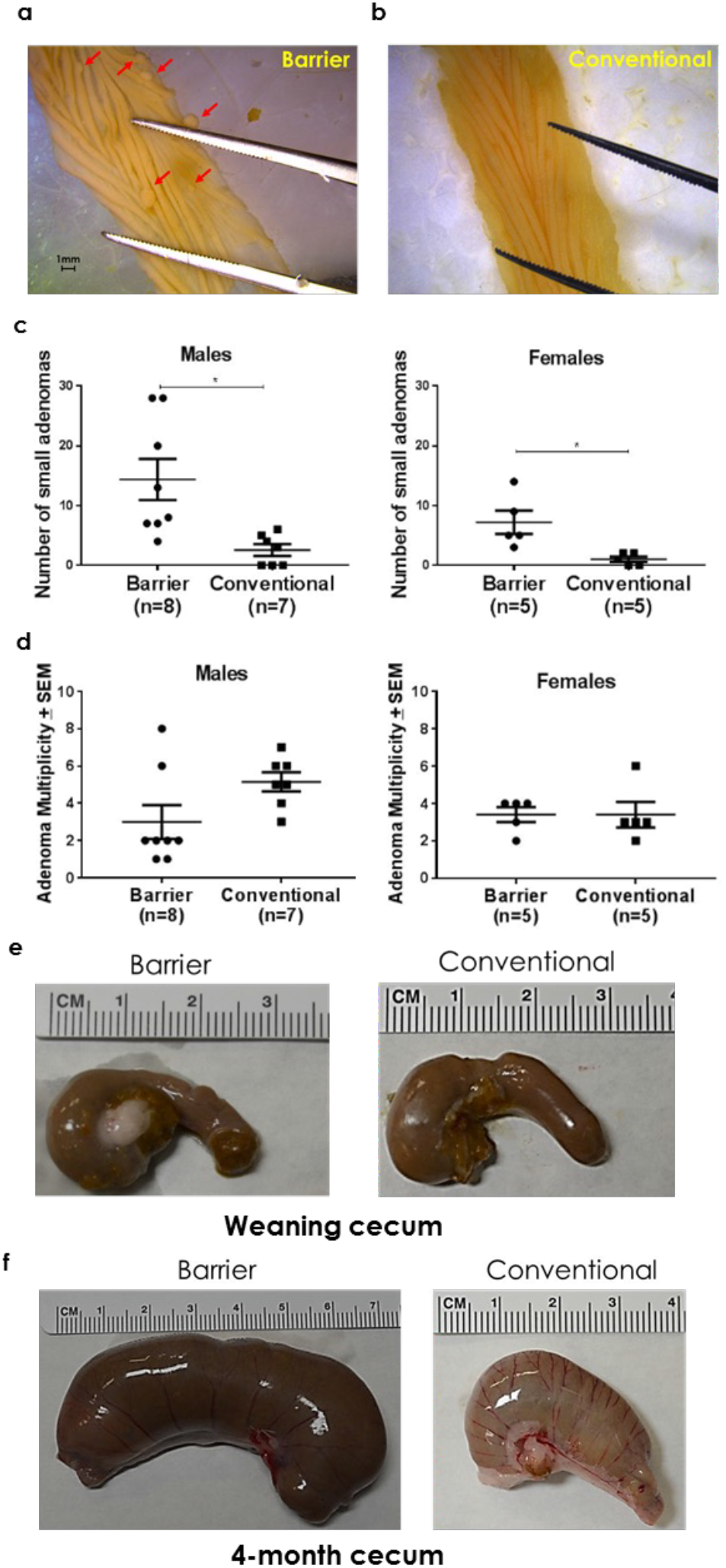
Effect of Altered Schaedler Flora on the colonic adenoma phenotype and the physiology of the gastrointestinal tract. (a) Representative proximal colon section of the (n=13) F1 Pirc rats from the barrier room. Arrows indicate small adenomas, less than 1 mm in diameter. Scale bar = 1mm. Depicted small adenoma sizes: 1 = 0.363mm, 2 = 0.858mm, 3 = 0.875 mm, 4 = 0.993mm, 5 = 0.969mm, and 6 = 0.378mm. (b) Representative proximal colonic region for (n=12) conventionally-housed rats. Images were captured on a Leica M165FC microscope with 1X magnification and a 40X objective. (c) Number of small adenomas determined in males and females respectively in the barrier and conventional rooms. (d) Adenoma multiplicity differences in males and females respectively were determined by excluding the number of small adenomas seen in the F1-Pirc rats. Significance was assessed by a Student’s *t-*test, with a *p-*value less than 0.05 was observed as significant. Error bars indicate standard error of the mean (±SEM). (e) Representative images of the cecum at weaning, from the barrier and conventional rooms. (f) Barrier and conventional room ceca obtained at sacrifice (representative images), indicating the difference in size between the housing conditions. Images were captured using a Nikon D5200. A ruler is shown for comparison between groups.

**Table 1:** Altered Schaedler Flora alters the colonic adenoma phenotype and the physiology of the gastrointestinal tract. Number of adenomas found in the small intestine (SI) and colon. Colonic adenomas smaller than 1 mm of diameter, in addition to the colonic adenomas are also provided. The last column indicates the presence or absence of small, proximal colonic adenomas which are not typically seen in the Pirc model of colon cancer.

| Animal<br># | Sex | Room | SI<br>adenomas | Colonic<br>adenomas<br>(diameter<br>more than<br>1mm) | Small<br>colonic<br>adenomas<br>(diameter<br>less than<br>1mm) | Proximal<br>small<br>adenomas |
| --- | --- | --- | --- | --- | --- | --- |
| 1852 | F | Barrier | 5 | 3 | 9 | Yes |
| 1853 | F | Barrier | 3 | 4 | 0 | No |
| 1882 | F | Barrier | 3 | 2 | 14 | Yes |
| 1902 | F | Barrier | 8 | 4 | 5 | No |
| 1920 | F | Barrier | 2 | 1 | 5 | Yes |
| 1863 | M | Barrier | 12 | 6 | 7 | Yes |
| 1883 | M | Barrier | 11 | 2 | 20 | Yes |
| 1884 | M | Barrier | 14 | 1 | 28 | Yes |
| 1886 | M | Barrier | 10 | 1 | 28 | Yes |
| 1908 | M | Barrier | 21 | 8 | 4 | No |
| 1926 | M | Barrier | 11 | 2 | 13 | Yes |
| 1940 | M | Barrier | 11 | 2 | 8 | Yes |
| 1943 | M | Barrier | 14 | 2 | 7 | Yes |
| 1860 | F | Conventional | 9 | 3 | 0 | No |
| 1879 | F | Conventional | 6 | 2 | 2 | No |
| 1880 | F | Conventional | 5 | 6 | 2 | No |
| 1922 | F | Conventional | 4 | 3 | 1 | No |
| 1934 | F | Conventional | 5 | 3 | 0 | No |
| 1865 | M | Conventional | 23 | 5 | 4 | Yes |
| 1887 | M | Conventional | 17 | 3 | 3 | No |
| 1890 | M | Conventional | 18 | 6 | 6 | No |
| 1891 | M | Conventional | 14 | 4 | 5 | No |
| 1927 | M | Conventional | 20 | 6 | 0 | No |
| 1930 | M | Conventional | 24 | 7 | 0 | No |
| 1938 | M | Conventional | 16 | 5 | 0 | No |

Furthermore, we sacrificed a cohort of F1 Pirc ASF animals at weaning and found no differences in their cecal size (Fig.3e). However, sacrifice after housing under barrier or conventional settings for 4 months, revealed considerable differences in cecum size. We found that rats maintained in the barrier room had ceca that were nearly 2-fold larger compared to conventionally-housed rats (Fig.3f). These results suggest that the lack of taxa from the conventional GM and/or their interactions with the Altered Schaedler Flora (ASF) in barrier rats is capable of modifying the physiology and the phenotype of the F1 Pirc rats.

### Conventional housing affects the GM architecture at 4 months of age

Considering the husbandry, handling, cecal and tumor multiplicity differences between the barrier and conventional rooms, we used 16S rRNA gene sequencing to determine the GM architecture in the F1-Pirc rats. At weaning, we found that the GM of rats at the time of separation into barrier and conventional rooms were similar to each other as indicated by bar graph (Fig.4a) and the Principal Coordinate analysis in Fig.4b (using Bray-Curtis distance matrix) and the overall richness determined by the number of OTUs observed in the samples (Fig.4c). They also resembled the GM profile of the parents, except for the conspicuous decrease in the relative abundance of genus *Mucispirillum* (Fig.4a).

**Figure 4.**
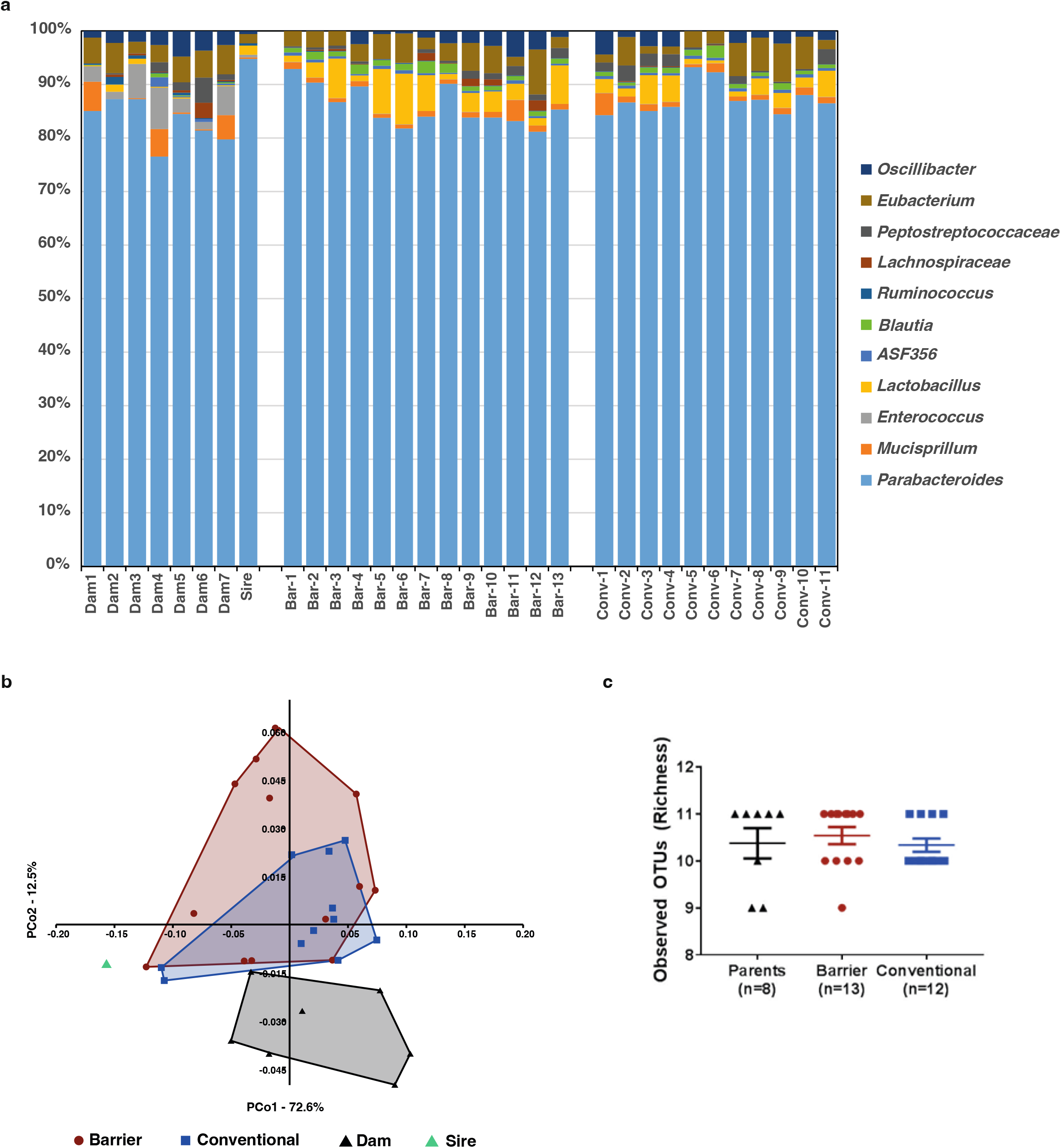
16S sequencing analysis of fecal microbiota in F1-Pirc rats at weaning. (a) Gut microbiota profiles of the dams and sire along with the barrier and conventionally raised F1-Pirc rats at weaning are displayed as a bar graph depicting the relative abundance of each OTU in percentages. Each color represents a single OTU. (b) Principal Coordinate Analysis using a Bray-Curtis distance matrix depicts the overall similarity or dissimilarity within the groups: barrier (brown, filled circle), conventional (blue, filled square), dams (black, filled triangle), and sire (green, filled triangle). PERMANOVA was used for significance testing; F=1.112 and *p*=0.3172. (c) The total number of OTUs observed, i.e. richness of the groups is depicted with the groups along the x-axis and the number of OTUs along the y-axis. No significant differences were found (ANOVA, Tukey’s post hoc, *p*<0.05)

At sacrifice (4 months of age), considerable differences were observed in the overall Genus’ profile of the GM between the barrier and conventionally-housed F1-Pirc rats (Fig.5a). At the genus level (Supplementary Fig.2A), several taxa including *Parabacteroides, ASF356, Blautia, and Mucispirillum* were elevated in the barrier F1-Pirc rats. In the conventionalized rats there was an observed increase in the relative abundance of over 50 taxa, the top 35 are depicted in the heatmap (Supplementary Fig.2a). The overall GM profile composition differences are visualized using a Principal Coordinate analysis (Fig.5b). The most separation was observed along PCo1, suggesting that the room differences contribute to the majority of the variability in the GM architecture. There were also significant increases in the richness and diversity indices such as Chao1 and Shannon (Fig.5c-e). These results suggest that the contribution of the room differences, such as husbandry, handling, and exposure to conventional animals have a crucial effect on the acquired taxa. *Firmicutes* and *Tenericutes* were increased in the conventionally-housed rats, whereas *Bacteroidetes* was decreased. This also led to a significant shift in the *Firmicutes:Bacteroidetes* ratio between the two groups (Fig.5f).

**Figure 5.**
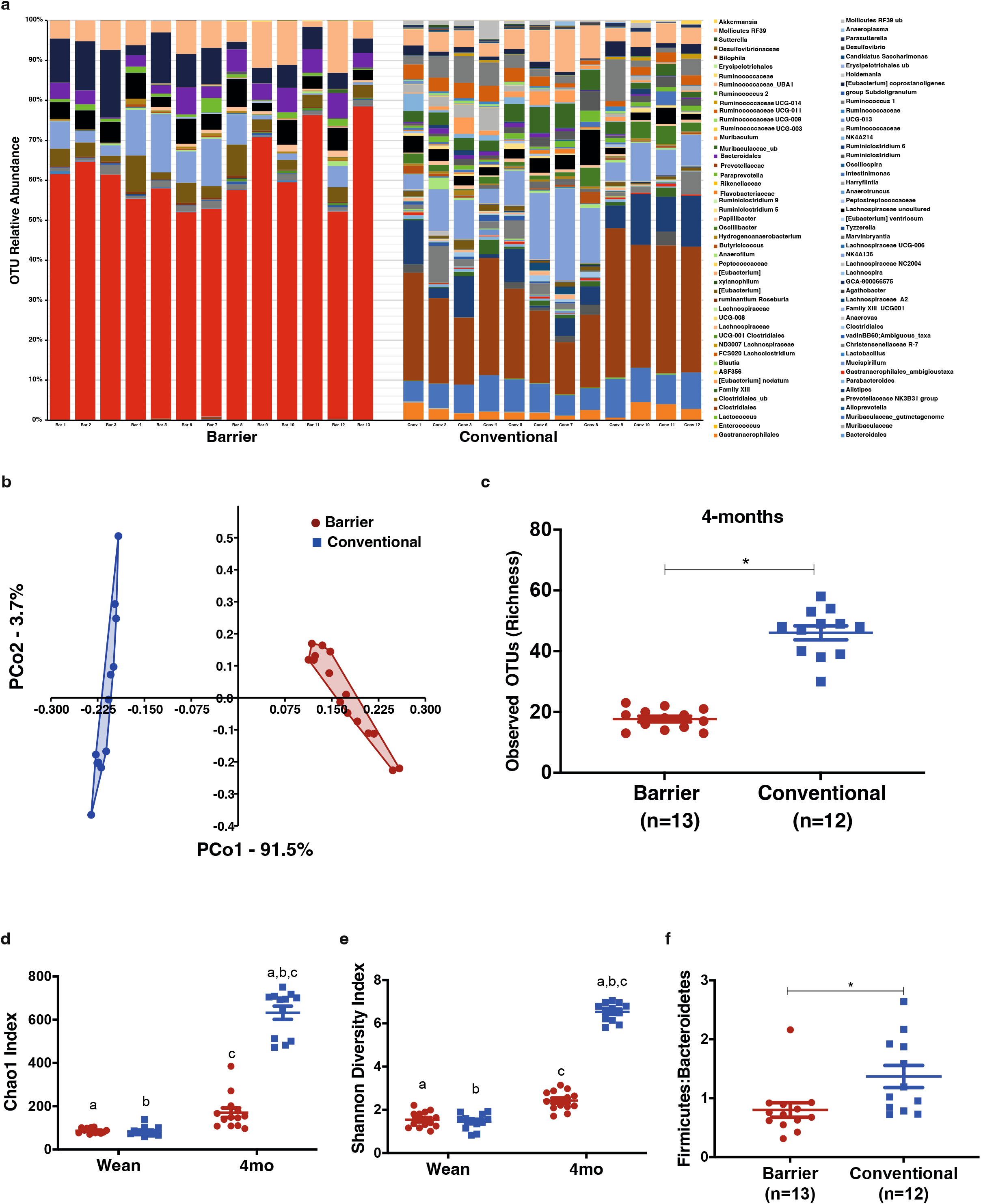
Effect of conventional housing on the GM at 4 months of age. (a) Bar graphs depicting the Phyla and Genus at 4-months of age from the barrier and conventional F1-Pirc rats demonstrate the individual OTUs as a different color. (b) Genus level OTUs were used to visualize the similarities/dissimilarities between each samples and the groups at 4-months of age using a PCoA. PERMANOVA was used to determine significant differences with a *p-*value less than 0.05. Based on the Genus data with a cutoff of 0.001% (accounting for sequencing error rates), the richness (c), and diversity indices – Chao1 (d) and Shannon (d) were measured from the raw read counts after normalizing the sequences to 21,639 per sample. (f) The *Firmicutes:Bacteroidetes* ratio of the two housing strategies is depicted. Significance assessed by *p*<0.05 was determined using a Student’s t-test.

We found significant correlations between certain taxa from the barrier (Fig.6a and Supplementary Fig.3a) room at weaning with the colonic tumor burden including the small adenomas. In these F1-Pirc rats, decrease in *Erysipelotrichaceae* and the genus *Parabacteroides* were associated with an increase in the colonic tumor count, whereas order *Peptostreptococcaceae* was found to show a positive correlation with tumor burden. Other taxa such as *Ruminococcaceae* and *Lachnospiraceae* showed similar correlations. Similarly, *Bacteroides, Peptococcus, Clostridiales, Peptococcaceae* and *Candidatus saccharomonas* showed significantly positive correlations.

**Figure 6.**
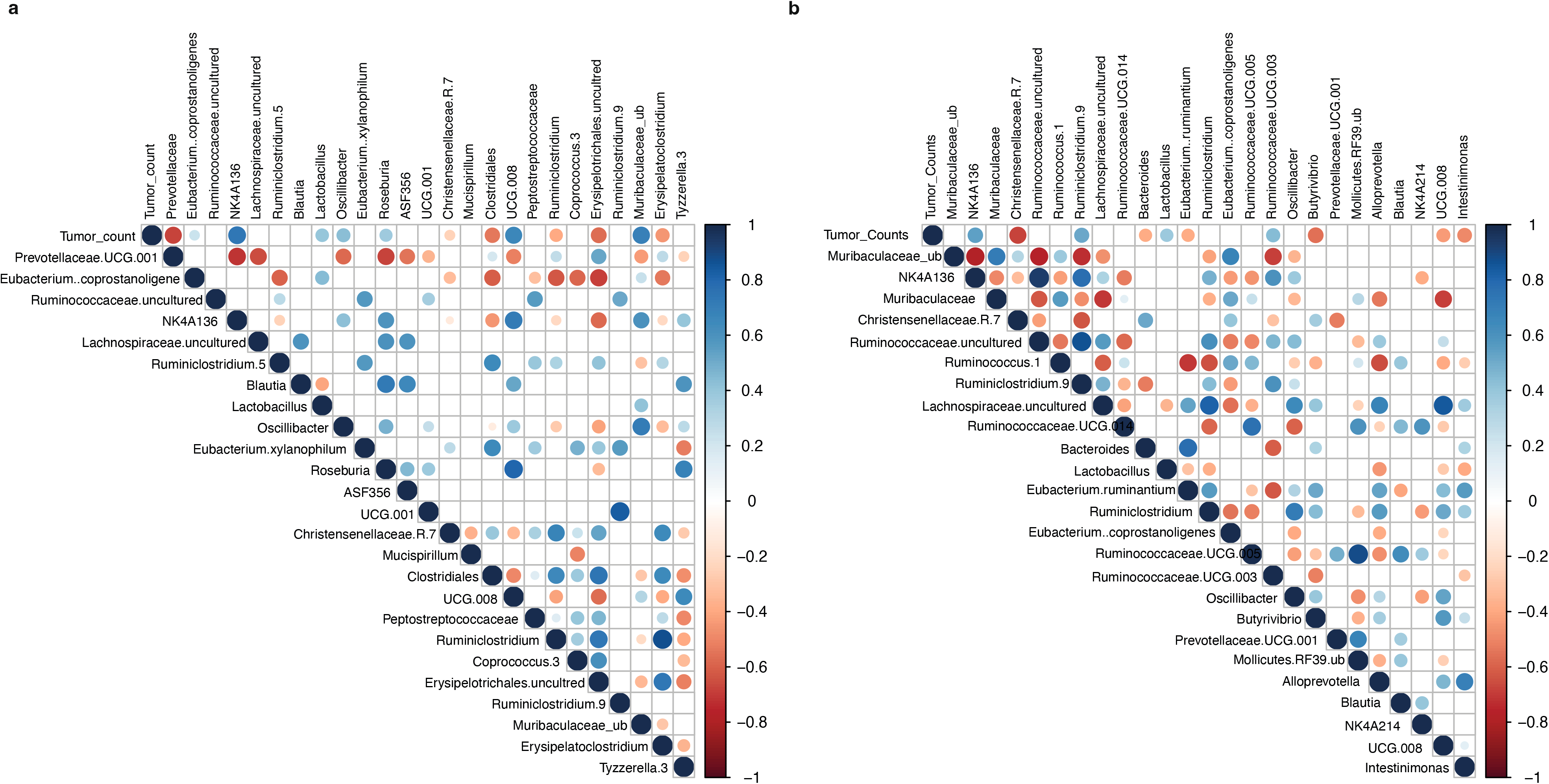
Correlation analysis of OTUs from barrier and conventional rooms with colonic tumor count at 4 months of age. Correlation analyses was performed using *Corrgram* R package, to determine positive or negative correlations with individual taxa at weaning in the barrier (a) and conventional (b) housing conditions. Correlations with a significant *p*-value of less than 0.05 are depicted by filled circles or diamonds. Empty cells indicate no significant correlations. Positive correlations with r^2^>0.75 are shown as blue diamonds and as blue circles for r^2^<0.75. Negative correlations are shown as red diamonds (r^2^>0.75), and red circles for <0.75.

Correlation analysis with the colonic tumor burden of the conventionally raised F1-Pirc animals (Fig.6b and Supplementary Fig.3b) showed that family *Prevotellaceae* at weaning was negatively correlated with tumor burden. *Clostridum Family XIII, Lachnospiraceae, Ruminococcus UCG-006, UCG-008, UCG-010* and *Clostridiales*, on the other hand were positively correlated with tumor increase, suggesting a causative role for the taxa from these operational taxonomic units. Simultaneously, we found significant negative correlations between *Parabacteroides* and specific OTUs from the barrier room at 4 months of age. These OTUs included *Ruminococcus NK4A136, Roseburia, Lachnospiraceae, Instestimonas and Oscillibacter*. Other OTUs including *Parabacteroides* and *Prevotellaceae* had positive and negative correlations respectively with the other commensals such as *ASF356, Mucispirillum, Lachnospiraceae* and *Ruminococcus UCG* taxa.

We used PICRUSt (Phylogenetic Investigation of Communities by Reconstruction of Unobserved States) and the HMP Unified Metabolic Analysis Network (HUMAnN) to understand the functional capacity of bacterial taxa in the fecal samples collected at 4 months of age (Supplementary Fig.2b). We found that the barrier room F1 Pirc rats had a substantial increase in the Spermidine-putrescine transport system and the succinate dehydrogenase pathways. Conversely, the predicted functional capacity of the conventional room rats showed increased abundance of pathways related to bacterial cell doubling time including Kreb’s cycle, increased amino acid biosynthesis (methionine and leucine), the iron transport system and increased sulfate reduction to hydrogen sulfide.

### Barrier and conventional room diets have distinct GM populations

Targeting the 16S rRNA gene, we sequenced the feed from the barrier and conventional rooms where we found that the GM profile of the feed from the two rooms did not differ significantly (Supplementary Fig.4a). Feed from the barrier and conventional rooms demonstrated similar abundances of order *Streptophyta* and *Zea luxurians*; genetic content likely derived from plant material used in the preparative process for feed (Supplementary Fig.5a). Though the community profile appeared similar, the relative abundances of *Lactobacillus, Leuconostoc, Sphingomonas* and *Fusobacterium* were significantly increased in the barrier room feed compared to the conventional diet (Supplementary Fig.5c). More importantly, to delineate between residual genetic content after autoclaving and potential taxa that may colonize the rats in the conventional room we cultured the feed from both rooms. Under anaerobic conditions, we cultured the feed overnight at 37 °C using brain heart infusion media. We observed several taxa in the feed from the conventional room grew abundantly, whereas the barrier room chow had minimally detectable levels of taxa such as *Clostridium* and *Bacillus* (Supplementary Fig.4b). This was also observed in the rarefaction curves (Supplementary Fig.5b) when sampling the observed species in each of the samples. We found that the cultures with the conventional feed had several species that were identifiable compared to both the uncultured conventional and barrier feed, including the cultured barrier feed. This suggested that bacteria from the feed could potentially have colonized the gastrointestinal (GI) tract of the F1 Pirc rats that were housed under conventional conditions, possibly altering the phenotype. In fact, we found that operational taxonomic units (OTUs) found in the feed such as *Lachospiraceae, RF39, Ruminiclostridium, Oscillibacter, Ruminicoccaceae* and several genus’ of the *Muribaculaceae* order were found in the conventionally raised F1-Pirc animals, but were undetectable in the barrier room rats (Supplementary Fig.5a).

## Discussion

The human colon is host to approximately 10^14^ bacteria alone, aside from viruses and fungi, which together form the gut microbiota. The interaction between the host and the endogenous GM is highly varied and complex which may be a crucial part of disease susceptibility. However, modelling the interactions of the GM in a complex setting is challenging. Therefore, we generated F344/NTac-*Apc^+/am1137^* (Pirc) rats and fostered them onto a Charles River Altered Schaedler Flora (CRASF) gut microbiota profile. We were able to stably maintain the ASF GM with only the acquisition of a few OTUs such as *Anaerotruncus, and Staphylococcus* etc. More importantly, the F1-Pirc (LEWF344F1-*Apc^+/am1137^* CRASF) animals generated resembled the CRASF parents at weaning except genus *Mucispirillum* which was decreased in the offspring compared to the breeders. This taxon is difficult to culture *in vitro* compared to other ASF taxa, however, it is still not known whether it is inhibited by the presence of other OTUs usually observed in altered Schaedler flora colonies ^14^. Based on reports of the CRASF populations by Matthias *et al ^15^* we find a majority of the original CRASF populations within our colony.

Dove et al ^16^ showed that germ-free model of the Apc mice developed two-fold fewer adenomas than the conventional. In rats, Reddy et al ^17^ also showed that germ-free animals, though treated with chemical carcinogens, had fewer tumors than conventional rats. Similarly, previous studies by Zackular *et al* ^3^, where the same model was treated with antibiotics showed a signficiant reduction in both small intestinal and colonic tumors. We hypothesized based on the phenotype of colon cancer in germ-free or antibiotic-treated animal models, that the F1-Pirc CRASF rats in the barrier room would have fewer colonic adenomas. Contrary to our hypothesis, animals maintained under barrier conditions had an increased tumor burden, including significantly increased number of smaller adenomas especially in the proximal section of the colon. In the original report of the Pirc rat ^2,3^, microadenomas required histopathological confirmation and were recorded as being smaller than 0.5 mm in diameter. However, in our case the differences between the barrier and conventional rooms were grossly apparent, including an increase in the number of smaller adenomas when compared to previous reports of conventional and germ-free rats. This forms an interesting hypothesis as to why and which OTUs of the GM may be initiating the shift from a distal colonic to a proximal. This disparity will be addressed in future studies. It must be noted though that excluding the smaller adenoma numbers, there was a trend towards increased tumors in the conventional animals as originally hypothesized. This posits for future studies where F1-Pirc rats would be aged longer than 4 months to understand if the observed small adenomas may develop into adenomas larger than 1 mm in diameter. Another observation from our study was the alteration of the colonic tumor phenotype observed in the F1-Pirc rats from the barrier room. Typically, Pirc rats demonstrate a colonic phenotype where the adenomas develop in the middle and distal portion of the colon with few in the proximal regions, as we observed in the F1-Pirc animals from the conventional room. However, F1-Pirc animals from the barrier room had several adenomas in the proximal section of the colon with few or none in the middle and distal regions. Similar to previous reports of germ free animals, these animals had enlarged ceca compared to conventional F1-Pirc rats ^18,19,20.^ Zackular *et al*. showed in an AOM/DSS-treated mouse model that a decrease in the overall GM population through the administration of antibiotics, led to a significant decrease in tumor burden. Another study similarly demonstrated that transferring tumor-associated microbiota into germ-free mice increased the tumor burden of the mice, otherwise significantly reduced when mice were maintained germ-free ^21^. Based on these reports, our findings of animals maintained in a barrier room having significantly elevated adenomas is intriguing. Although studies have shown that bacteria are needed for a phenotype to be manifested in animals ^22,23^, our results suggest that a consortium of taxa may influence disease. Also, our observations that a simple, yet diverse consortium of taxa within the gut microbiome, still maintains a large cecal size as compared to conventionally-housed rats is notable. However, in previous reports by ^19,23,24^, the cecal size is reported as a measure of percent body weight. Future studies will analyze the differences in cecal sizes compared to other germ-free or mono-colonized models. In 1962, Skelly et al ^25^ reported that certain Clostridium species restored cecal size to what is observed in normal conventional settings. Alternatively, Loesche in 1969 reported that B.fragilis caused an increase in cecal size ^24,25^. We observed both Clostridium and Bacteroides species within our microbiome analyses, and will need additional studies including germ-free, ASF, and conventional Pirc rats to identify the source of this variation.

The barrier room was maintained with irradiated chow, paper chip bedding, autoclaved water and animals were always handled in a biosafety cabinet. We housed the conventional room rats with non-irradiated feed, non-autoclaved bedding and used animal handling techniques that did not require aseptic methods. We hypothesized that this would alter the existing CRASF microbiota to a more complex GM. We used 16S rRNA sequencing to determine if the GM, known to be modulated by husbandry factors ^2626–28^ was the crucial modulator of the phenotype observed in our study at 3-months after introduction into the conventional facility, we found the conventional rats had acquired OTUs including *Prevotellaceae, Ruminococcaceae, Muribaculaceae, Parasutterella* and *Desulfovibrionaceae*. *Prevotellaceae* and *Desulfovibrionaceae* have been reported to be associated with healthy patients or a decreased tumor burden in colon cancer studies ^3,10,29^. On the other hand, *Blautia, Enterococcus*, and some *Lachnospiraceae* taxa found in the barrier room F1-Pirc rats have been associated with an increased tumor susceptibility ^30–33^. This was equally evident from the correlations where *Peptococcaceae, Clostridiales, Lachnospiraceae*, previously reported to be associated with an increased tumor burden were elevated and positively correlated with the tumor burden in the barrier rats ^34,35^. Correlation analysis also found that certain OTUs introduced into the conventional rats had a negative association with *Parabacteroides*, potentially suggesting that these OTUs inhibit the proliferation or take over the niche occupied by the latter, i.e. competitive interactions ^36^. In the barrier room rats we also found predicted functional pathways such as succinate dehydrogenase and spermidine-putrescine transport system to be elevated. Host succinate dehydrogenase mutations are very commonly found in colon cancer ^37,38^. This raises the possibility of a breakdown of the host dehydrogenases, thereby leading to an increase in the bacterial dehydrogenase expression to counteract the toxic effect of succinate. Alternatively, many rumen bacteria are known to produce succinate which in turn has been identified as a biomarker for colon cancer via mass spectrometry ^39^. This suggests elevated levels of succinate, reportedly an onco-metabolite ^40^ could be promoting tumorigenesis in the barrier room rats via inhibition of PHD (prolyl hydroxylase domain-containing) enzymes ^41^ via activation of hypoxia-induced factor alpha (HIF-α). Succinate quantitation via metabolomics and PHD enzyme activity will however need to be validated in future studies to determine the mechanisms contributing to increased succinate levels. Similarly, polyamines such as spermidine and putrescine have been reported to be biomarkers for colorectal cancer in human patients ^42^. In 1988, Upp *et al*. analyzed the polyamine levels including spermidine and putrescine in colon cancer patients and found that they may be used to identify at-risk patients of the disease ^42,43^. More recently, it was identified that GI bacteria such as *Bacteroides fragilis* upregulates spermine oxidase which induces production of spermidine, hydrogen peroxide and aldehydes ^42–44^, potentially causing DNA damage. Another thought-provoking observation in our study is the presence of OTUs found in the diet that were detected in the barrier and conventional room fecal samples from F1-Pirc rats. Although, the barrier room rats were not over-ridden by the taxa found via 16S sequencing, this was not true for the conventionally raised rats. We found that the conventionally housed F1-Pirc rats had significant amounts of bacterial taxa that were also detected in the diet, and that were anaerobically cultivable. This suggests that the non-irradiated diet, may be one source of the variation, although it is also possible that the rare OTUs picked up are nonviable residual DNA from dead bacteria or spores. More importantly, this source, potentially led to a significant shift in the phenotype, i.e. number of adenomas.

Colorectal cancer (CRC) animal models have been extensively used to study and understand the etiology of the disease including initiation, development and factors affecting susceptibility. Despite the development of the *Apc^+/am1137^* rat, the *Apc^+/Min^* mouse model of colon cancer is still largely used for various studies owing to cost and the ease of genetic manipulation techniques. However, the Pirc (*Apc^+/am1137^*) rat with a colonic phenotype has created a potentially more translatable alternative to the mouse when studying colon cancer. While not the primary focus of this study, we realize that it is not inconceivable that there may be inherent differences between the gut microbiome of wild type and CRASF-Pirc rats and that genetics play a crucial role in the composition of the GM. However, with studies recently reporting evidence of the role of the gut microbiota in diseases susceptibility including colon cancer ^3,4,6,33^, the importance of reproducibility in disease models is critical. Several reports have identified *Fusobacterium*, in particular *Fusobacterium nucleatum*, as a significant modifier of disease burden ^45,46^. This bacteria along with Enterotoxigenic *Bacteroides fragilis* (ETBF) has often been associated with increased tumor burden and/or carcinoma samples in human patients ^4,7^. However, it should be noted that most of these studies do not take into account the constant interactions and synergistic nature of the commensals within the GI tract. GM populations are a constant source of nutrients and metabolites, which are contingent on the action of one bacterium on the by-products of the replicative processes of another. To model and establish a simplified GM profile to study the role of specific bacteria and their interactions with the host and other commensals, we established *Apc^+/am1137^* rats on a CRASF (Charles River Altered Schaedler Flora) gut microbiome profile. The observance of increased tumor number in a limited GM microbiome provides a platform for probiotic experimentation. It can also allow for more refined metabolite profiling and longitudinal assessment in changes in metabolic processing. Utilizing a simplified GM profile for understanding the pathophysiology of colon cancer, may provide insights into the interactions between commensals and with the host, including the mechanisms by which specific taxa promote or prevent adenomagenesis.

## Methods

### Animal Care and Use

All procedures were performed according to the guidelines regulated by the Guide for the Use and Care of Laboratory Animals, the Public Health Service Policy on Humane Care and Use of Laboratory Animals, and the Guidelines for the Welfare of Animals in Experimental Neoplasia and were approved by the University of Missouri Institutional Animal Care and Use Committee.

### Charles River Altered Schaedler Flora (CRASF) rats and cross-fostering

7 week old Lewis rats with a limited Altered Schaedler Flora (n = 4 males, and 4 females) were purchased from Charles River Laboratories Inc. Laboratories (Wilmington, MA). The animals were shipped overnight in a sterile double-enclosed isolator cage with sterile bedding and Hydrogel® gel packs (Portland, ME) to the Discovery Ridge animal facility at University of Missouri. Fecal samples were collected prior to shipping and upon arrival at the facility for 16S rRNA sequencing. Simultaneously, bedding and gel-pak samples that the animals were shipped with were also collected for sequencing. The animals were housed in a barrier room on ventilated racks (Thoren, Hazleton, PA) in micro-isolator cages with autoclaved paper chip bedding (Shepherd Specialty Paper, Milford, NJ) and were fed irradiated 5053 PicoLab Mouse Diet 20 (LabDiet, St. Louis, MO), and water (reverse osmosis purified and sulfuric acid (pH 2.5-2.8) acidified followed by autoclaving), and allowed to acclimatize for a week, after which they were setup into breeder pairs. Timed matings for fostering were set up with our F344/NTac *Apc^+/am1137^* (generation, N=28) conventional rat colony.

Female F344/NTac rats were checked for plugs, and on day 21 post observation of plugs, a Caesarean was performed. The uterus was tied-off at both ends prior to surgical resection and then transferred in a sterile petri dish with betadine solution to the barrier room. In a biosafety hood, the uterus was opened with a pair of sterile scissors and the pups were physically manipulated after removing the amniotic sac and warmed under a heat lamp. Only CRASF breeders with pups on the ground within 36 hours were used as surrogates for fostering the F344/NTac-*Apc^+/am1137^* pups. Half the litter and bedding was removed from the CRASF surrogate, and mixed with the to-be fostered pups thoroughly, before placing the F344/NTac-*Apc^+/am1137^* fostered pups along with a few of the CRASF pups with the surrogate mom. At 12 days of age, all pups including the fostered ones were clipped for genotyping.

### Genotyping and animal identification

Pups were ear-punched prior to weaning at 12 days of age using sterile technique. DNA was extracted using the “HotSHOT” genomic DNA preparation method previously outlined ^47^. DNA was used for genotyping using a high resolution melt (HRM) analysis as described previously ^10,47^.

### Experimental design, animal husbandry (breeding) and barrier room housing

F1-Pirc rats were generated by crossing one founder male, F344/NTac-*Apc^+/am1137^* CRASF Pirc rat established via cross-fostering, with wild-type female LEW/Crl ASF rats. The rats were housed on ventilated racks (Thoren, Hazleton, PA) in micro-isolator cages as described above. Animal handling required complete personal protective equipment (PPE) including face masks, hair nets and TyVek sterile sleeves (Cat.No.17988110, Fisher Scientific, USA) and the use of 10% bleach spray to treat surfaces including gloves and sleaves. Prior to breeding fecal samples were collected from both the breeders using aseptic methods. LEWF344F1-*Apc^+/am1137^* (F1 generation) ASF pups were generated and genotyped at 12 days of age.

### Conventional room housing

At weaning, F1 Pirc rats were co-housed in the conventional room with F344/NTac animals from the holding colony with an endogenous complex GM when available, in microisolator cages on ventilated racks with nonsterile paper chip bedding. Cage changes for conventional rats were done on open benches. Rats in the conventional room were fed non-irradiated 5008 Lab diet and had *ad libitum* access to acidified (sulfuric acid, pH 2.5-2.8), autoclaved water.

### Fecal sample collection

Fecal samples were collected from the pups at weaning, and monthly thereafter starting until sacrifice at 4-months of age. Briefly, fecal samples were collected by placing the animal in a clean, sterile cage without bedding. Freshly evacuated feces were speared with a sterile toothpick or forceps and placed into a sterile Eppendorf tube. All samples were stored at −80 °C until further processing.

### Fecal DNA extraction, 16S library preparation, sequencing and analysis

Fecal samples were pared down to 65 mg using a sterile blade and then extracted using methods described previously ^10^. Amplification and sequencing of the V4 hypervariable region of the 16s rDNA was performed at the University of Missouri Metagenomics center and DNA core facility (Columbia, MO) and the results annotated using the SILVA 16S database ^10,48^. Samples with a read count below 15,000 were removed from the analysis due to insufficient rarefaction. The average read counts for all samples was 57,863. Microbial Community DNA Standards from ZymoBIOMICS ™ were used to account for any errors via extraction and sequencing processes.

All OTUs with a relative abundance below 0.001% were excluded from analysis. Principal Coordinate analyses were performed in PAST (PAleontological STatisitcs, version 3.2) ^49^. PERMANOVA with default permutations (N=9999) was used to determine significant differences between groups when performing PCoA analyses using the module embedded into PAST3.2. Simultaneously, a scree plot was generated using the chemometrics.R script under the m*etaboanalyst* package to identify which principal coordinates to plot for the figures. Heatmaps were generated using the plotHeatMap function from the same package along with the hclust function from the *stat* package. For the heatmaps, Euclidean distance was used as the similarity measure, while Ward’s clustering algorithm accounting for average linkage was used to create the Dendrogram. Correlation analyses testing the relationship of OTUs’ relative abundance with tumor burden was assessed using the *corrgram* package in R (version 3.4.1), assessing the top 50 OTUs based on the individual relative abundance. PICRUSt, HUMAnN and LEfSE analysis was performed after re-annotating (closed-reference) the 16S rDNA gene sequences against the Greengenes (May, 2013) database as described previously ^49,50^.

### Anaerobic culturing of the lab diet feed and DNA extraction

3 samples of 0.5g of feed from the barrier and conventional rooms was introduced anaerobically into an autoclaved serum vial, closed with a sterile rubber stopper and an aluminum crimp seal. Oxygen was purged from vials and 5ml of sterile brain heart infusion (BD Difco, ThermoFisher Scientific, USA) media was added using a syringe. The inoculum and media was then incubated anaerobically overnight at 37 °C in a 5% CO2 incubator. After incubation, the contents of the vial were used for DNA extraction using previously established methods including manual DNA precipitation and the DNEasy kit (Qiagen, USA) ^51^.

### Tumor counts

All animals were humanely euthanized with CO_2_, administration and necropsied at 16 weeks of age. The small intestine and colon from the rats were placed on to bibulous paper and then splayed opened longitudinally. Tissues were then fixed overnight in Carnoys fixative (30% Choloroform, 10% glacial acetic acid and 60% absolute ethanol), and were replaced with 70% ethanol for long term storage until adenoma counting was performed.

### Statistical analyses and figures

Statistical analyses and graphing for figures (except Fig.1) were prepared through GraphPad Prism version 7 for Windows (GraphPad Software, La Jolla, CA). *p*-values were set to identify significance at a value less than 0.05, unless otherwise indicated.

### Ethics Approval and Consent to Participate

The current study was conducted in accordance with the guidelines set forth by the Guide for the Use and Care of Laboratory Animals and the Public Health Service Policy on Humane Care and Use of Laboratory Animals. All studies and protocols were approved by the University of Missouri Institutional Animal Care and Use Committees.

## Availability of Data and Material

The sequences and raw 16S rDNA read files generated and analyzed during the current study are available at the NCBI GenBank database under BioProject ID# PRJNA490292.

## Acknowledgements

The authors wish to acknowledge Giedre Turner, Becky Dorfmeyer and the MU Metagenomics Center (MUMC) for their assistance with 16S rRNA gene sequencing; Brittany Lister and Office of Animal Resources staff for assistance with animal husbandry; Charles River Laboratories Inc. for assistance making the CRASF animals available and with sample collection prior to shipping. This research was funded by the University of Missouri System Research Board grant. The funders had no role in study design, data collection and analysis, decision to publish, or preparation of the manuscript.

## Author Contributions

Conceived and designed the experiments: S.B.B., J.M.A.L., D.D. Performed the experiments: S.B.B. J.M.A.L., J.M. and D.D. Analyzed the data: S.B.B., D.D. and J.M.A.L, Manuscript preparation: S.B.B., J.M.A.L. All authors contributed to the review of the manuscript.

## Competing Interests

The authors declare no competing interests.

---

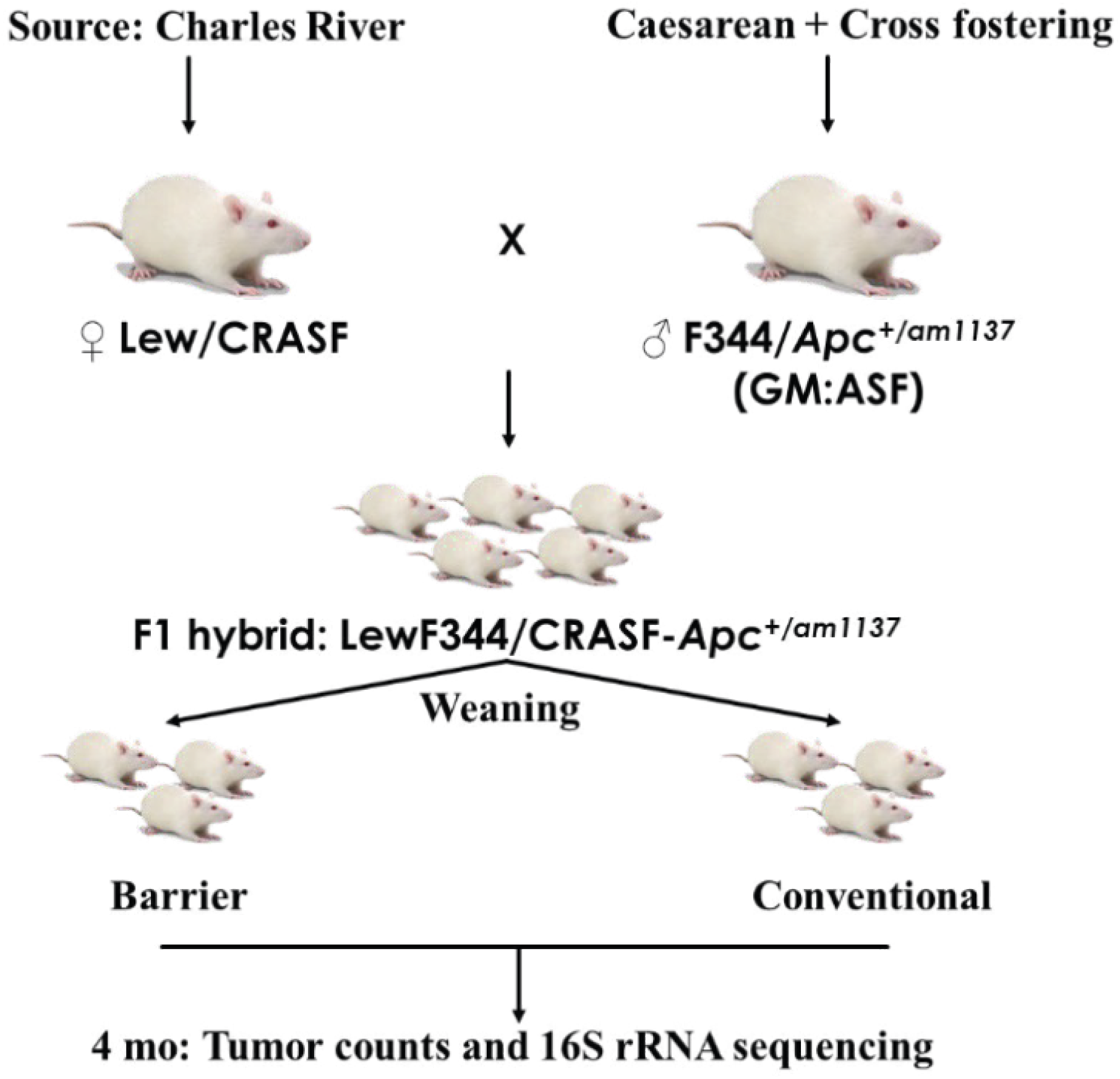

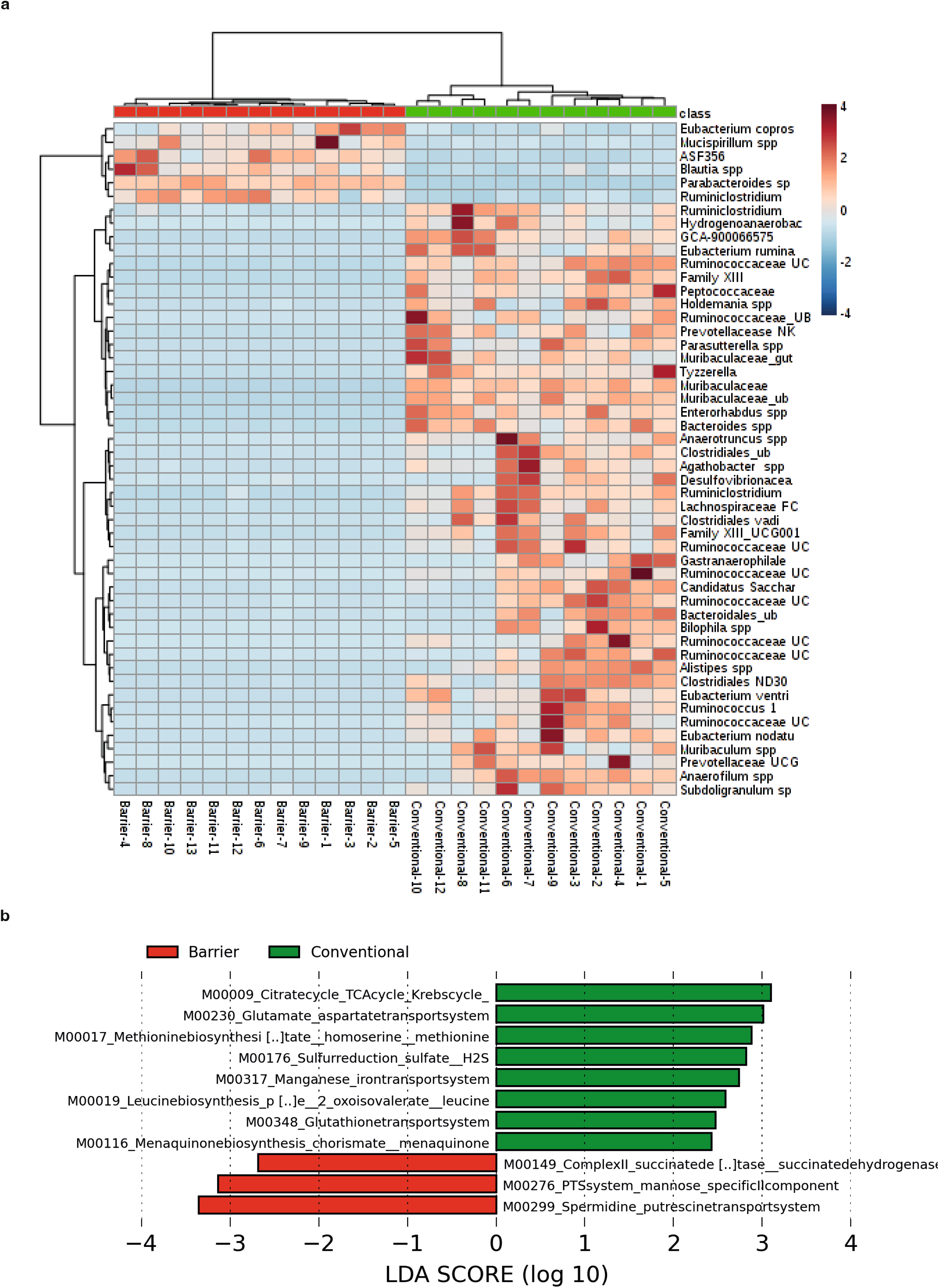

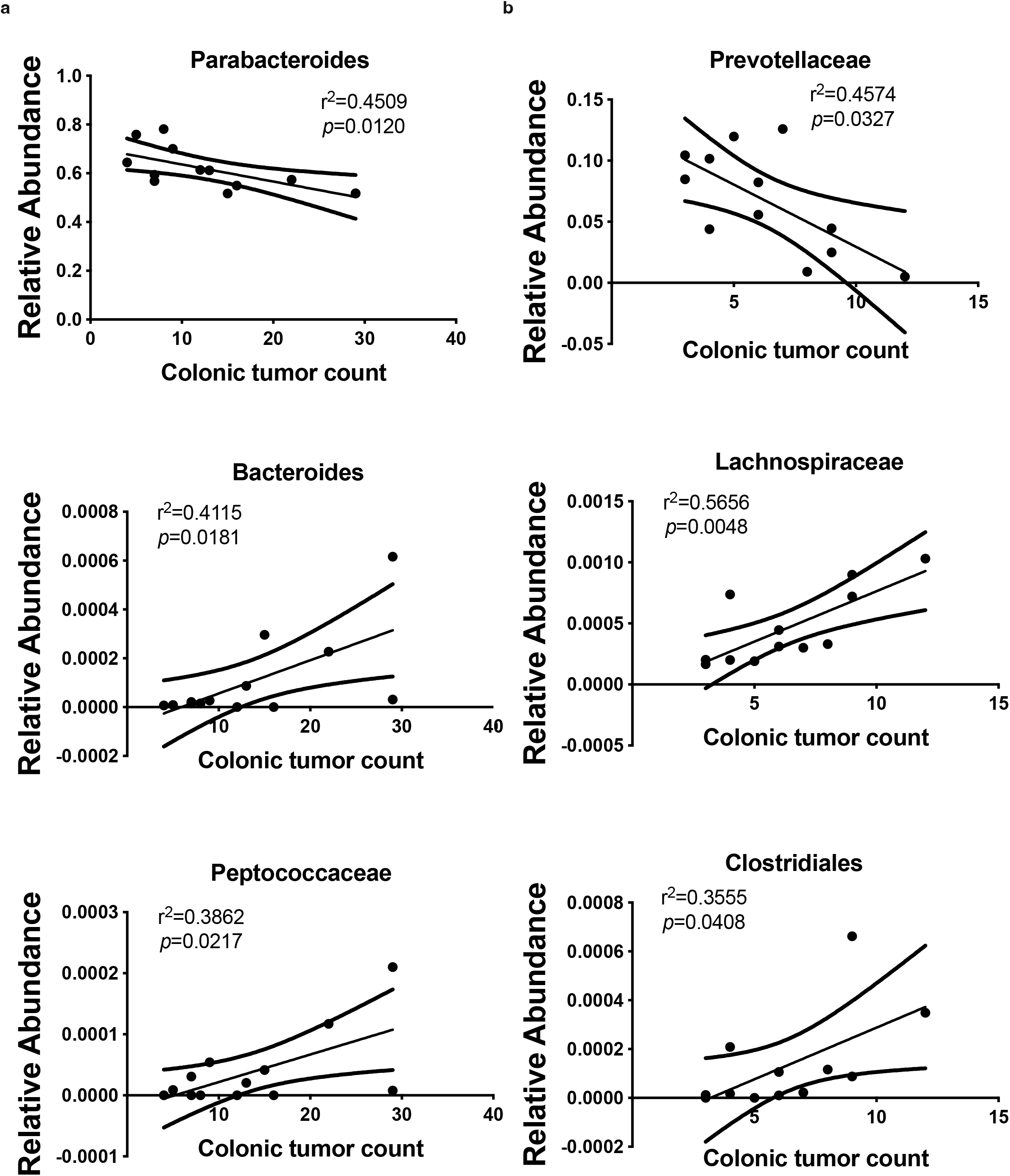

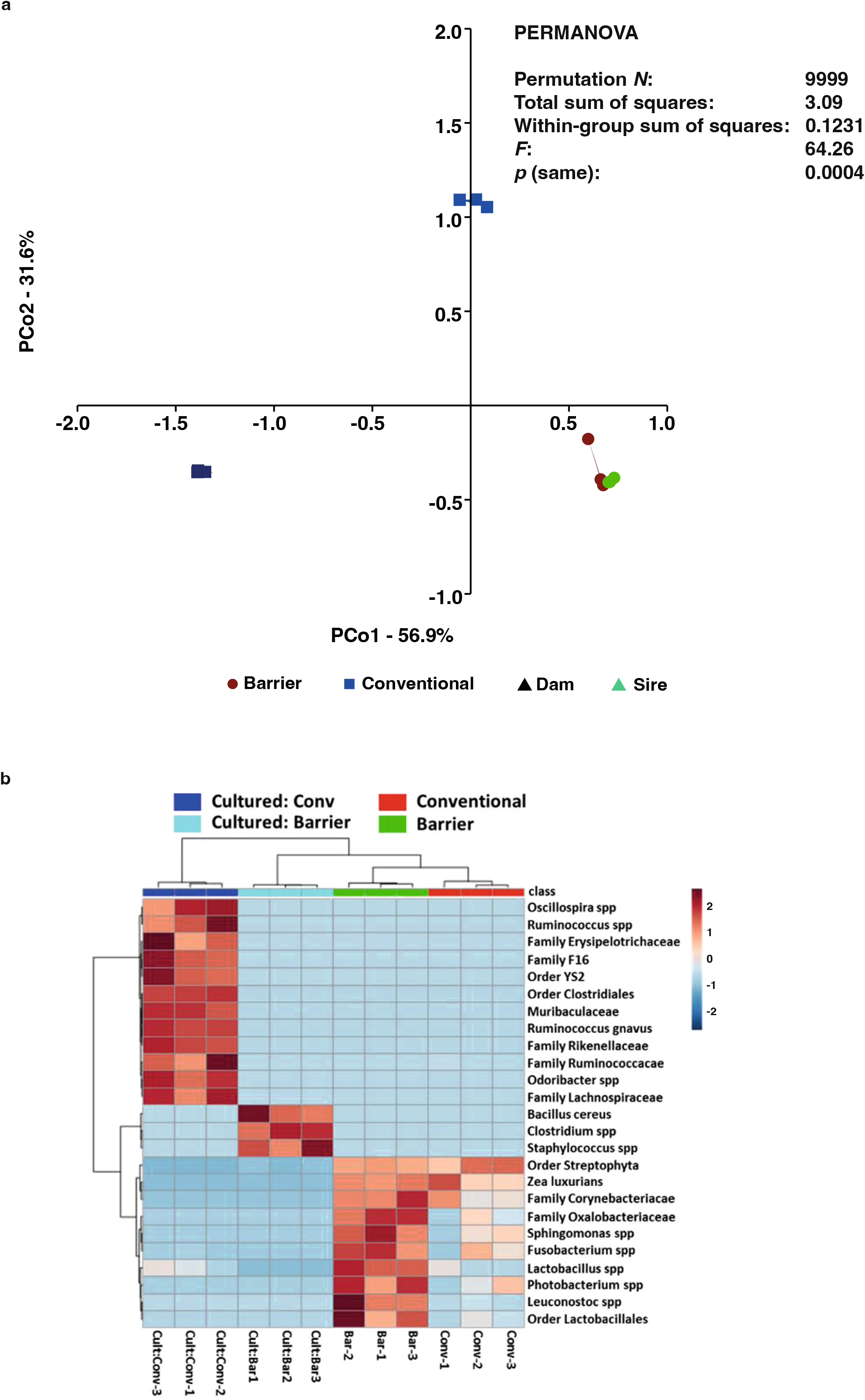

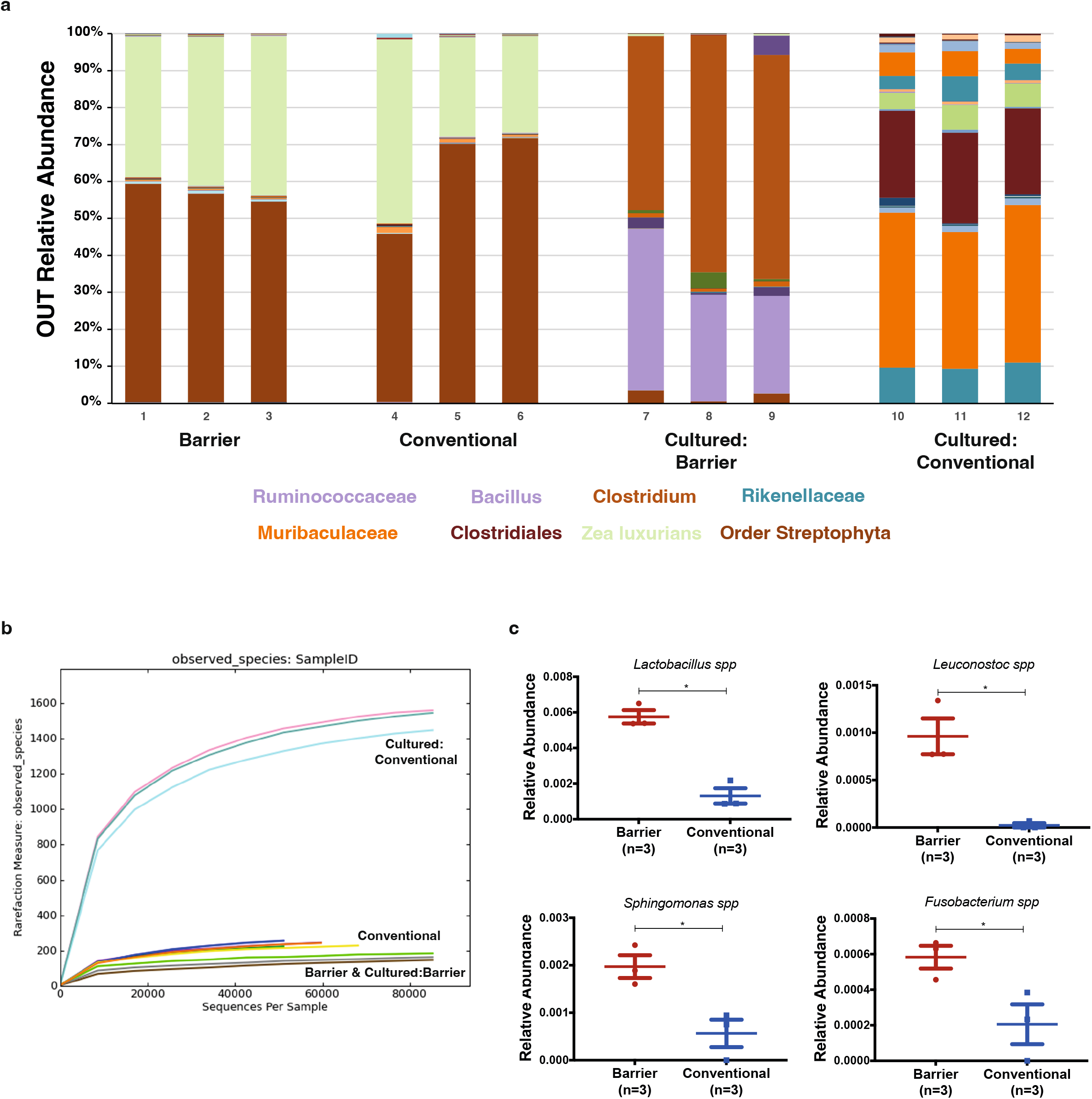

## Notes

### Competing Interest Statement

The authors have declared no competing interest.

### Summary of Updates

The revision of the manuscript include reference changes and an update of all figures due to poor quality initial figures.

