## Supplementary Figures for "Shift from a Simplified to Complex Gut Microbiota Reduces Adenoma Burden in a Preclinical Rat Model of Colon Cancer" (from another version of this preprint: version 1, 2023-01-23)

Supplementary figure 1. Experimental Design

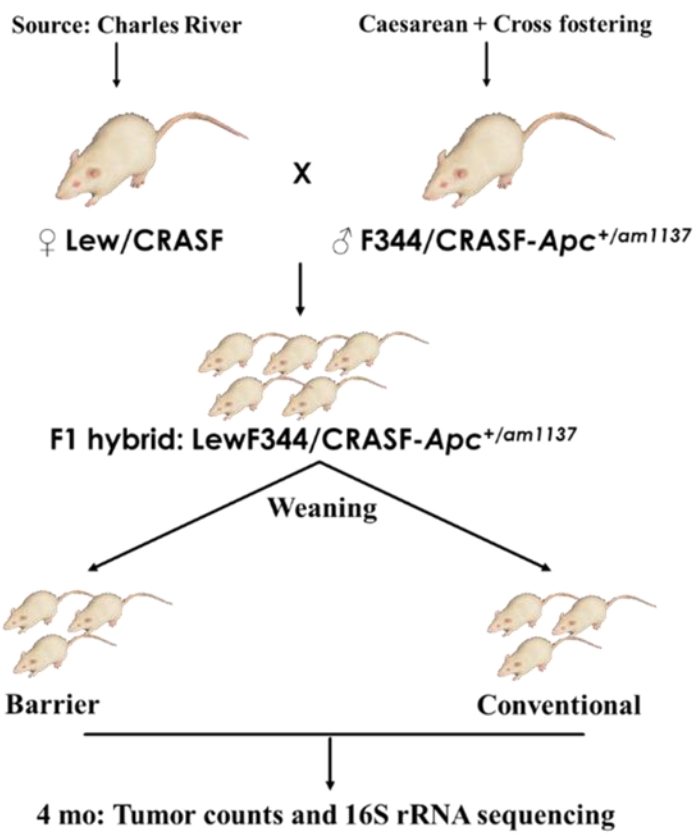


Supplementary figure 1. Experimental Design

Female LEW/CRASF rats were mated with male F344/CRASF-*Apc*$^{+/am1137}$ (Pirc) rats to generate F1-hybrid LEWF344F1-*Apc*$^{+/am1137}$ CRASF rats. At weaning, littermates were equally divided and housed either in a barrier room or in a conventional settings. At 4 months of age, colonic tumor counts and 16S rRNA sequencing was used to determine adenoma burden and gut microbiota (GM) profiles.

**Supplementary figure 2. GM profile and predicted metabolic function of barrier and conventional rats at 4 months of age**

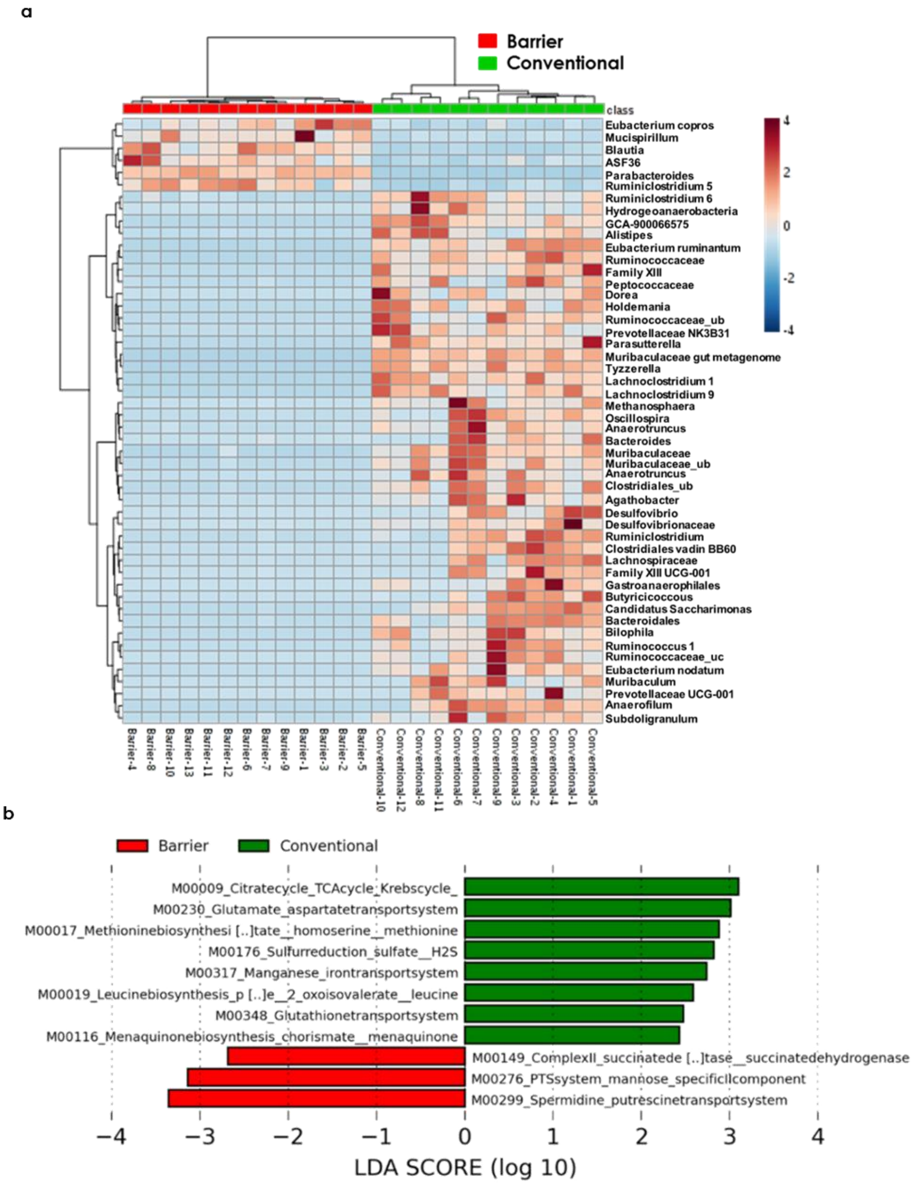


<u>Supp.fig.2. GM profile and predicted metabolic function of barrier and conventional rats at 4 months of age</u>

(a) Heatmap of the Genus' found at 4 months of age, generated using a Euclidean distance measure and Ward's clustering algorithm, depicts the top 50 OTUs that are differential between the groups. Barrier room animals are shown in red on the top bar, while the conventional animals are shown as green. (b) Linear Discriminant Effect Size (LEfSe) analysis of the predicted metagenomic pathways obtained via PICRUSt and HUMAnN between the conventional and barrier room animals was performed using fecal DNA on 16S rDNA gene sequencing at 4 months of age.

**Supplementary figure 3. Correlation analysis of OTUs with colonic tumor count**

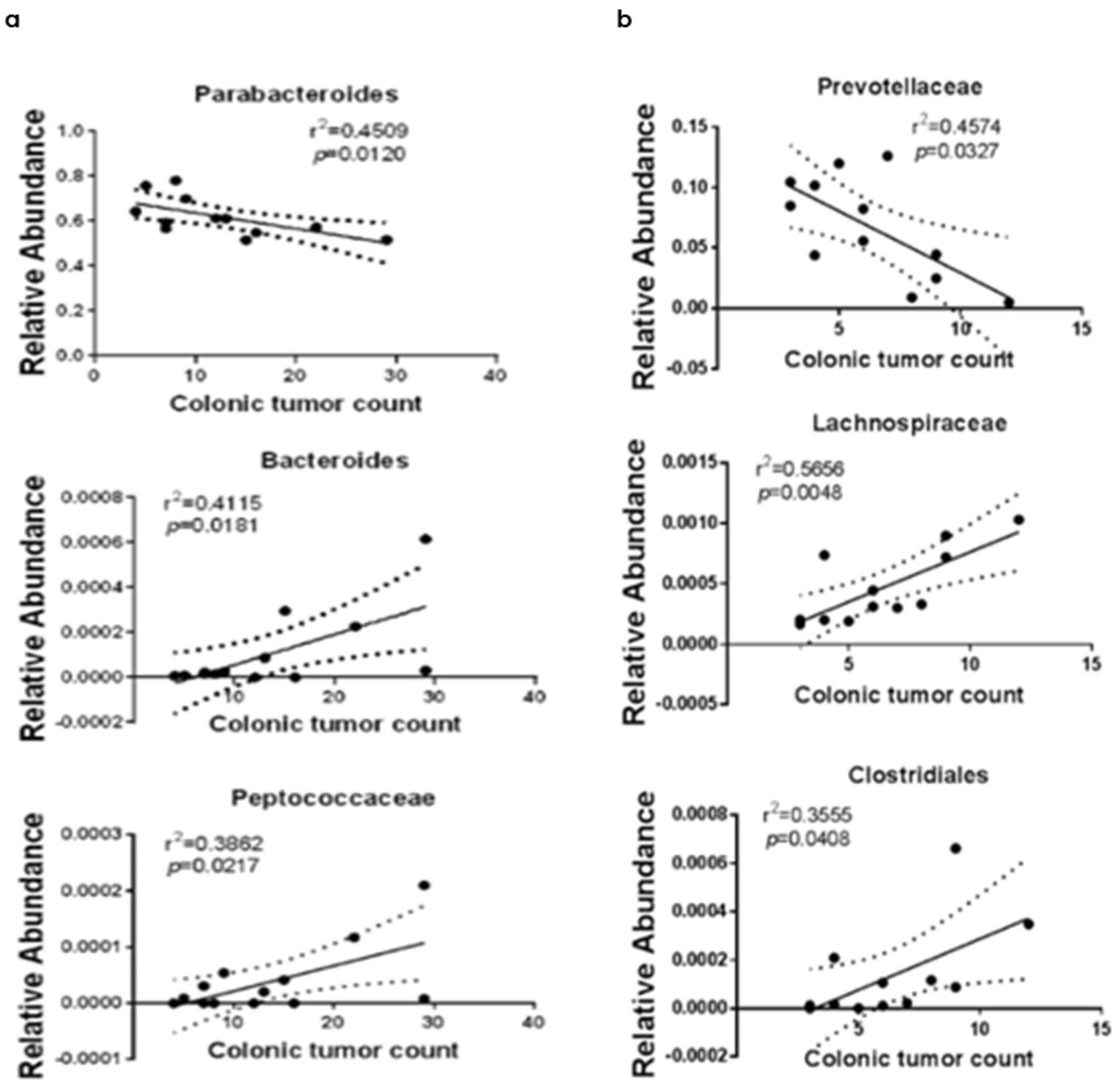


Supp.Fig.3. Correlation analysis of OTUs with colonic tumor count

Representative bacterial taxa relative abundance at 4 months of age correlated positively (*Bacteroides, Peptococcaceae, Lachnospiraceae* and *Clostridiales*) or negatively (*Parabacteroides* and *Prevotellaceae*) with colonic tumor count in the (a) barrier and (b) conventional F1-Pirc rats. Significance was determined using Pearson's correlation test with a $p<0.05$ considered to be significant.

**Supplementary figure 4. Bacterial population analysis of barrier and conventional room feed via 16S rDNA sequencing**

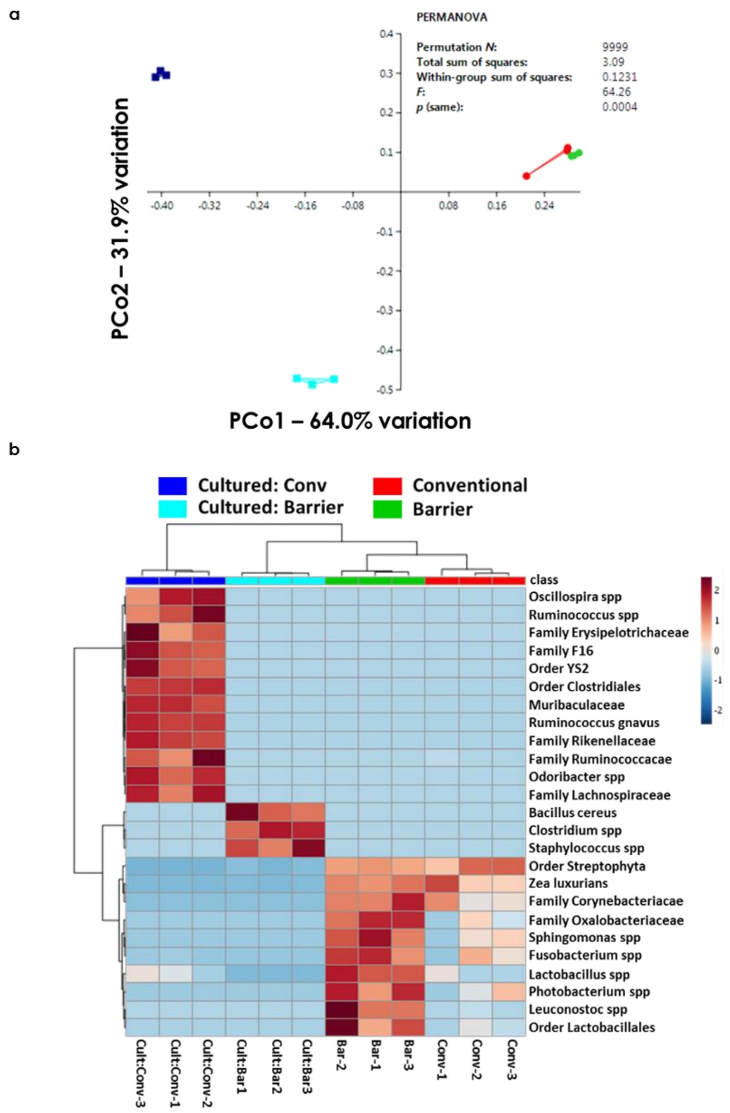


<u>Supplementary figure 4. Bacterial population analysis of barrier and conventional room feed via 16S rDNA sequencing</u>

(a) Principal Coordinate analysis (PCoA) demonstrates the overall GM profile variations along PCo1 and PCo2, between the feed from both the rooms. Each filled circle represents a single sample from the barrier (brown) and conventional (blue) room diet. No significant differences were found despite the separation via a PERMANOVA. (b) Heatmap generated using Euclidean distance and Ward's clustering algorithm was generated to observe the top 25 variable OTUs between the barrier (green) and conventional (red) feed 16S sequencing data. The Dendrogram clusters samples based on their similarity with each other, where the individuals OTUs are listed along the y-axis. Relative abundance of the OTUs are plotted in color with increased abundance indicated by the darker red and lower abundance highlighted by the darker blue shading.

### Supplementary figure 5. 16S analysis of cultured feed from barrier and conventional rooms

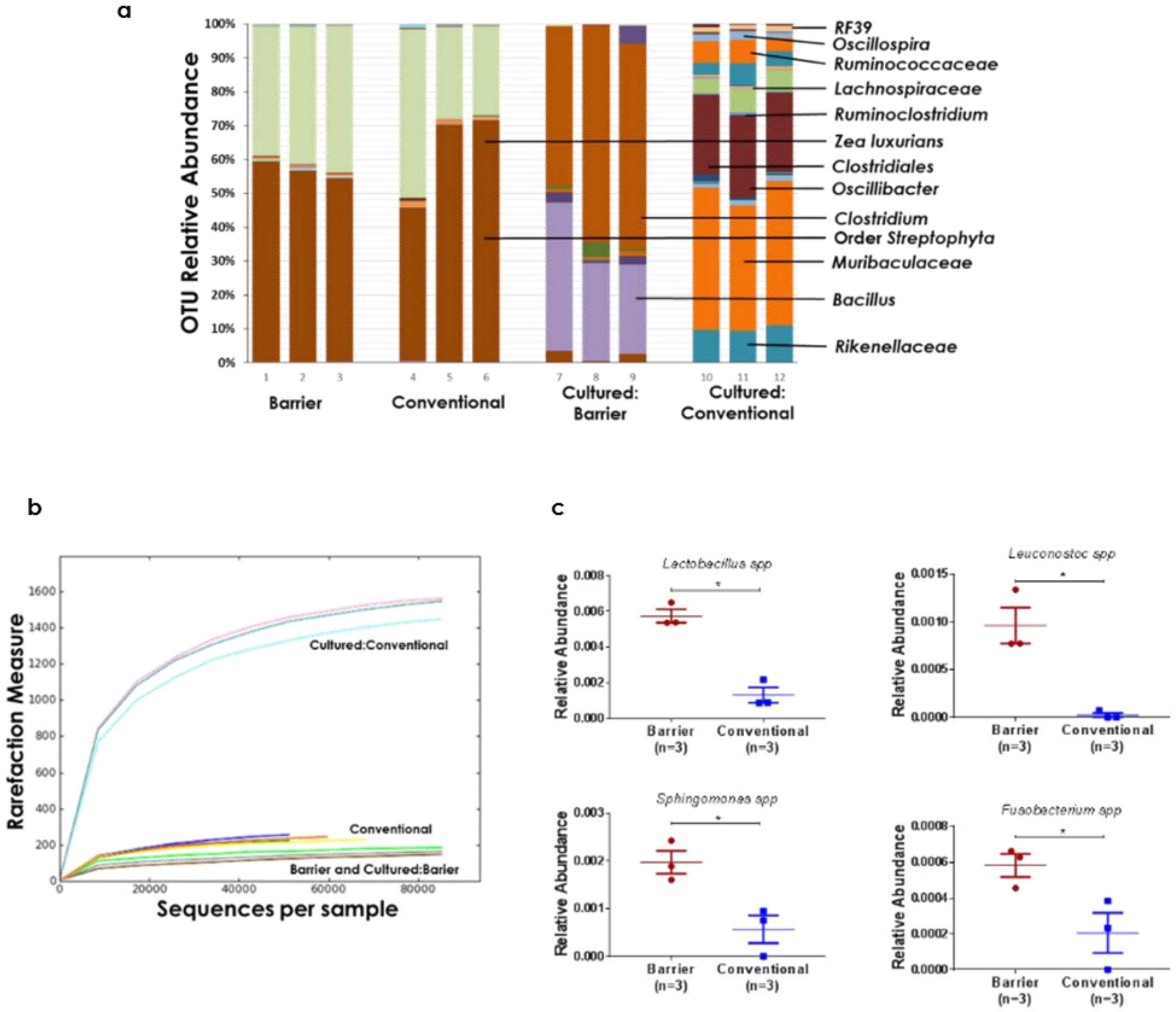


Supp.Fig.5. 16S analysis of cultured feed from barrier and conventional rooms

(a) 16S rRNA sequencing of the V4 hypervariable region from (n=3) diet samples is demonstrated as a stacked bar graph. Each color represents a single operational taxonomic unit (OTU) in the diet in terms of relative abundance along the y-axis. Groups depicted include: barrier diet, conventional diet, cultured barrier diet and cultured conventional diet. (b) Rarefaction curves displaying the depth of sequencing achieved for each sample in the barrier, conventional, and cultured barrier and conventional groups. Total number of observed species (y-axis) is plotted against the number of sequences (x-axis) per sample. (c) Relative abundance of individual bacteria found to be differential between the barrier and conventional feed. Significance was determined by a *p*-value of less than 0.05 using a Student's *t*-test.
